# Host infection by the grass-symbiotic fungus *Epichloë festucae* requires catalytically active H3K9 and H3K36 methyltransferases

**DOI:** 10.1101/2020.09.01.277384

**Authors:** Yonathan Lukito, Kate Lee, Nazanin Noorifar, Kimberly A. Green, David J. Winter, Arvina Ram, Tracy K. Hale, Tetsuya Chujo, Murray P. Cox, Linda J. Johnson, Barry Scott

## Abstract

Recent studies have identified key genes in *Epichloë festucae* that control the symbiotic interaction of this filamentous fungus with its grass host. Here we report on the identification of specific fungal genes that determine its ability to infect and colonize the host. Deletion of *setB*, which encodes a homolog of the H3K36 histone methyltransferase Set2/KMT3, specifically reduced histone H3K36 trimethylation and led to severe defects in colony growth and hyphal development. The *E. festucae* Δ*clrD* mutant, which lacks the gene encoding the homolog of the H3K9 methyltransferase KMT1, displays similar developmental defects. Both mutants are completely defective in their ability to infect the host grass, and mutational studies of key residues in the catalytic SET domains from these proteins show that these phenotypes are dependent on the methyltransferase activities of SetB and ClrD. A comparison of the differences in the host transcriptome between seedlings inoculated with wild-type versus mutants suggests that the inability of these mutants to infect the host was not due to an aberrant host defense response. Co-inoculation of either Δ*setB* or Δ*clrD* with the wild-type strain enables these mutants to colonize the host. However, successful colonization by the mutants resulted in death or stunting of the host plant. Transcriptome analysis at the early infection stage identified four fungal candidate genes, three of which encode small-secreted proteins, that are differentially regulated in these mutants compared to wild-type. Deletion of *crbA*, which encodes a putative carbohydrate binding protein, resulted in significantly reduced host infection rates by *E. festucae*.

**Author Summary:** The filamentous fungus *Epichloë festucae* is an endophyte that forms highly regulated symbiotic interactions with the perennial ryegrass. Proper maintenance of such interactions is known to involve several signalling pathways, but much less is understood about the infection capability of this fungus in the host. In this study, we uncovered two epigenetic marks and their respective histone methyltransferases that are required for *E. festucae* to infect perennial ryegrass. Null mutants of the histone H3 lysine 9 and lysine 36 methyltransferases are completely defective in colonizing the host intercellular space, and these defects are dependent on the methyltransferase activities of these enzymes. Importantly, we observed no evidence for increased host defense response to these mutants that can account for their non-infection. Rather, these infection defects can be rescued by the wild-type strain in co-inoculation experiments, suggesting that failure of the mutants to infect is due to altered expression of genes encoding infection factors that are under the control of the above epigenetic marks that can be supplied by the wild-type strain. Among genes differentially expressed in the mutants at the early infection stage is a putative small-secreted protein with a carbohydrate binding function, which deletion in *E. festucae* severely reduced infection efficiency.

## Introduction

Eukaryotic genomes are organised into discrete domains that are defined by the degree of chromatin condensation: euchromatin is generally less condensed and transcriptionally active, whereas heterochromatin is highly condensed and transcriptionally inactive [1]. The degree of chromatin condensation is controlled by various post-translational modifications of specific amino acid residues on the N-terminal tails of histones, including methylation, acetylation and phosphorylation [2, 3]. Cross-talk between the various protein complexes that modulate these histone modifications generates positive and negative feedback loops that control the final chromatin state and gene expression output [4, 5]. Among these various histone modifications, methylation of histone H3 is perhaps the most well-studied. Methylation of histone H3 lysine 4 (H3K4) and lysine 36 (H3K36) are associated with actively transcribed genes in euchromatic regions, whereas methylation of lysine 9 (H3K9) and lysine 27 (H3K27) is associated with the formation of constitutive and facultative heterochromatin, respectively [5]. These methyltransferase reactions are catalysed by SET domain proteins belonging to the COMPASS (K4), SET2 (K36), SpCLRC/NcDCDC (K9), and PRC2 (K27) protein complexes.

These histone marks and their respective methyltransferases are key regulators of fungal secondary metabolism [5–7], and are also required to establish and maintain the mutualistic interaction between the fungus *Epichloë festucae* and its grass host [8, 9]. Deletion of the *E. festucae* genes encoding the H3K9 methyltransferase ClrD or the H3K27 methyltransferase EzhB led to the derepression in culture of the subtelomeric *IDT* and *EAS* gene clusters, which encode the enzymes required for indole-diterpene (IDT) and ergot alkaloid (EAS) biosynthesis, respectively. Both mutations also impacted on the ability of *E. festucae* to form a symbiotic association with *L. perenne*; no host infection was observed after inoculation with the *clrD* mutant, whereas plants infected with the *ezhB* mutant had a late onset host phenotype characterized by an increase in both tiller number and root biomass [8]. Deletion of the *cclA* gene, which encodes a homolog of the Bre2 component of the COMPASS (Set1) complex responsible for deposition of H3K4me3, also led to transcriptional activation of the *IDT* and *EAS* genes in axenic culture. In contrast, deletion of the *kdmB* gene, which encodes the H3K4me3 demethylase, decreased *in planta* expression of the *IDT* and *EAS* genes, and reduced levels of IDTs *in planta* [10]. Plants infected with the *cclA* or *kdmB* mutants have a host interaction phenotype that is similar to wild-type, demonstrating that loss of the ability to add (Δ*cclA*) or remove (Δ*kdmB*) H3K4me3 does not impact on the ability of *E. festucae* to establish a symbiosis. To complete our analysis of the four major histone H3 methyltransferases we examine here the role of *E. festucae* SetB (KMT3/Set2) in the symbiotic interaction.

Set2 was originally identified in yeast as an enzyme that induces transcriptional repression through methylation of histone H3K36 in yeast [11]. The Set2 protein interacts with the hyperphosphorylated form of RNA polymerase II (RNAPII) and catalyses methylation of H3K36 during transcriptional elongation [12–14]. A role for H3K36 methylation in transcriptional repression has been shown for both yeast and metazoans [11, 15–17]. More recent studies showed that presence of H3K36me3 in filamentous fungi is not associated with gene activation [18, 19], and has an important role in regulating fungal development and pathogenicity. Deletion of *SET2* homologs in *Magnaporthe oryzae* [20], *Fusarium verticillioides* [21], and *Fusarium fujikuroi* [19] led to growth defects in culture and reduced host pathogenicity. Culture growth defects were also observed for the corresponding mutants in *Neurospora crassa* and *Aspergillus nidulans* [22–25] highlighting the importance of Set2 for fungal development. Here we show that a catalytically functional Set2 homolog is crucial for *E. festucae* to infect its host *L. perenne* and establish a mutualistic symbiotic association.

## Results

### SetB is an H3K36 methyltransferase that regulates fungal growth and development

The gene encoding the *E. festucae* homolog of the *Saccharomyces cerevisiae* H3K36 methyltransferase Set2 (KMT3) was identified by tBLASTn and named *setB* (gene model no. EfM3.042710; [26]). Protein structure analysis identified canonical PreSET, SET, PostSET, WW and SRI domains along with an additional TFS2N domain (Fig 1A), as reported for the *N. crassa* Set2 homolog [5]. *E. festucae setB* was deleted by targeted gene replacement using a *nptII* geneticin resistance cassette for selection. PCR screening of Gen^R^ transformants and Southern blot analysis identified a single Δ*setB* strain (S1 Fig). Western blot analysis of total histones showed that H3K36me3 was specifically depleted in this Δ*setB* strain, while levels of H3K36me1/2 were increased (Fig 1C). Introduction of the *setB* wild-type allele restored these methylation defects, confirming the role of SetB in H3K36 methylation (Fig 1C). This Δ*setB* strain grew extremely slowly in culture compared to the wild-type strain, a phenotype that was complemented by re-introduction of the *setB* wild-type allele (Fig 1D). Microscopic analysis of the hyphal morphology revealed that Δ*setB* hyphae had a wavy pattern of growth, branched more frequently and had hyphal compartments that were significantly shorter than observed for the wild-type strain (Fig 1E).

**Fig 1.**
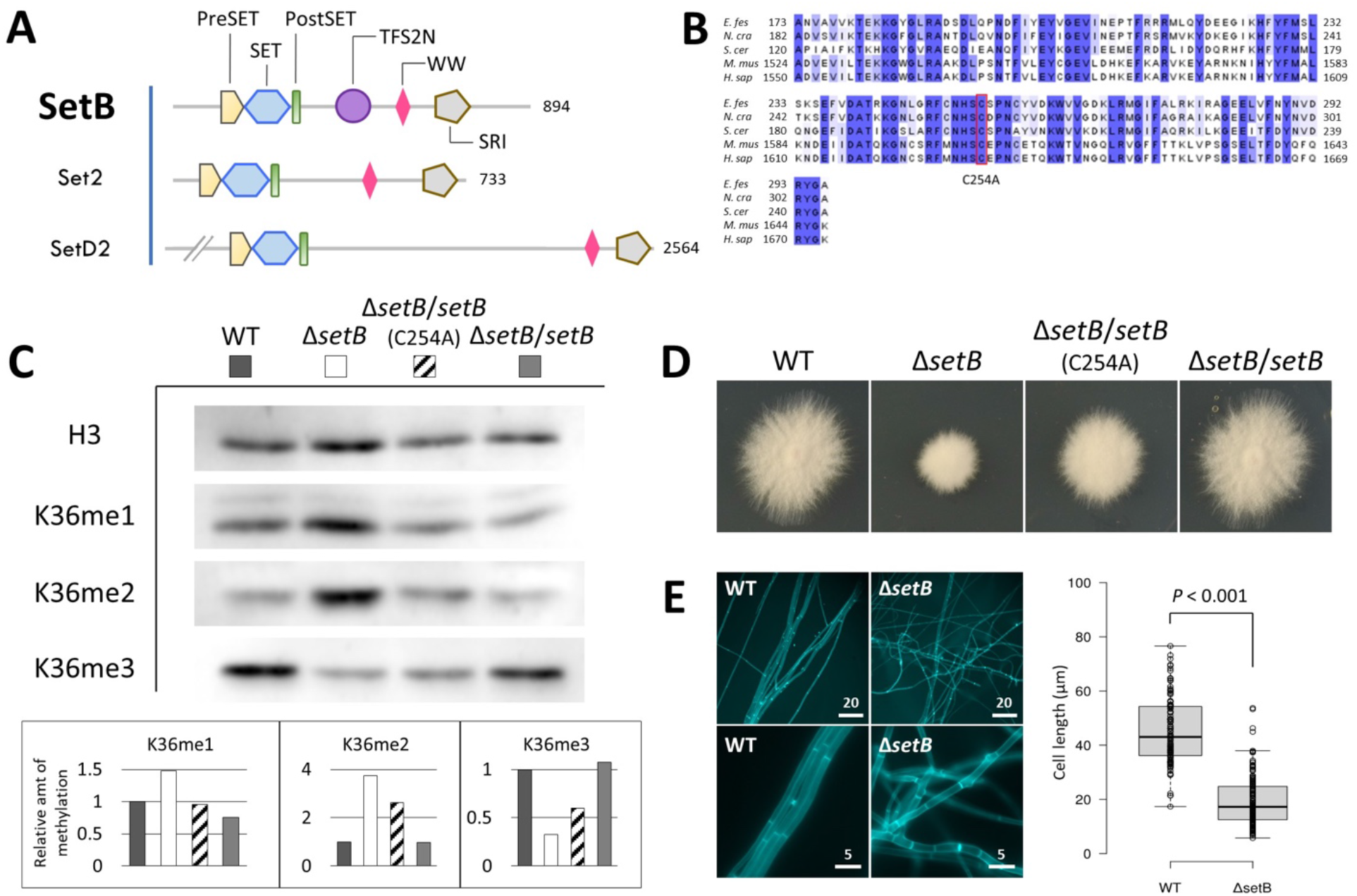
*E. festucae* SetB is an H3K36 methyltransferase. (A) Domain structure of *E. festucae* SetB and the yeast and mammalian orthologs. Numbers indicate the protein length in amino acids. (B) Amino acid alignment of SET domains of SetB and other SET2 proteins showing conservation of the cysteine 254 residue that was mutated to alanine in this study. (C) Western blot of total histones show reduced H3K36me3 and accumulation of H3K36me1/2 in Δ*setB*. Relative histone methylation was quantified by comparison to H3 band intensities, with the wild-type value arbitrarily set to 1. (D) Colony morphology after 6 days of growth in culture. (E) Deletion of *setB* affected hyphal morphology and length. Numbers above scale bars refer to length in μm. Boxplots represent more than 80 measurements of hyphal compartment lengths for each strain. The horizontal centre line of the boxes represent the medians and the top and bottom edges of the boxes represent the 75^th^ and 25^th^ percentiles, respectively.

### The H3K36 methyltransferase activity of SetB is required for *E. festucae* to infect *Lolium perenne*

To study the role of SetB in the symbiotic interaction of *E. festucae* with its host, *L. perenne* seedlings were inoculated with Δ*setB* and grown for 8-12 weeks, after which time mature plant tillers were tested for infection by an immunoblot assay using an antibody raised against *E. festucae*. Four independent inoculation experiments revealed that Δ*setB* was unable to infect these host plants (Table 1, Fig 2A). The absence of the mutant in these mature plants was confirmed by confocal laser-scanning microscopy (CLSM) (Fig 2B). To determine if Δ*setB* failed to infect and colonize leaf tissue, or whether it was lost from the leaf tissue during tiller growth (reduction of endophyte persistence), we also examined the infection status of seedlings at one- and two-weeks post inoculation (wpi) by CLSM. At both timepoints, both endophytic and epiphytic hyphae of wild-type were observed at the infection site, but only epiphytic and not endophytic hyphae of Δ*setB* (Fig 2C and D). These observations demonstrate that Δ*setB* is completely unable to colonize the host at the site of infection. Re-introduction of the wild-type *setB* allele into the mutant restored the ability of this strain to infect *L. perenne* (Fig 2B, Table 1).

**Table 1.** Host infection rates of *setB* and *clrD* mutants. Number of plants infected with the indicated strains, over the total number of plants in each independent study. Percentages of infection are indicated in parentheses. Plants were analysed at 8-12 wpi.

|  | **Wild-type** | **$\Delta setB$** | **$\Delta setB/setB$** | **$\Delta setB/setB^{(C254A)}$** |
| --- | --- | --- | --- | --- |
| Expt 1 | 20/37 (54%) | 0/44 (0%) | 18/27 (67%) | 10/38 (26%) |
| Expt 2 | 28/32 (88%) | 0/35 (0%) | 33/54 (61%) | 8/85 (9%) |
| Expt 3 | 33/46 (72%) | 0/36 (0%) |  |  |
| Expt 4 | 21/28 (75%) | 0/41 (0%) |  |  |
|  | **Wild-type** | **$\Delta clrD/clrD$** | **$\Delta clrD/clrD^{(C265A)}$** |  |
| Expt 1 | 10/15 (67%) | 12/13 (92%) | 0/52 (0%) |  |

**Fig 2.**
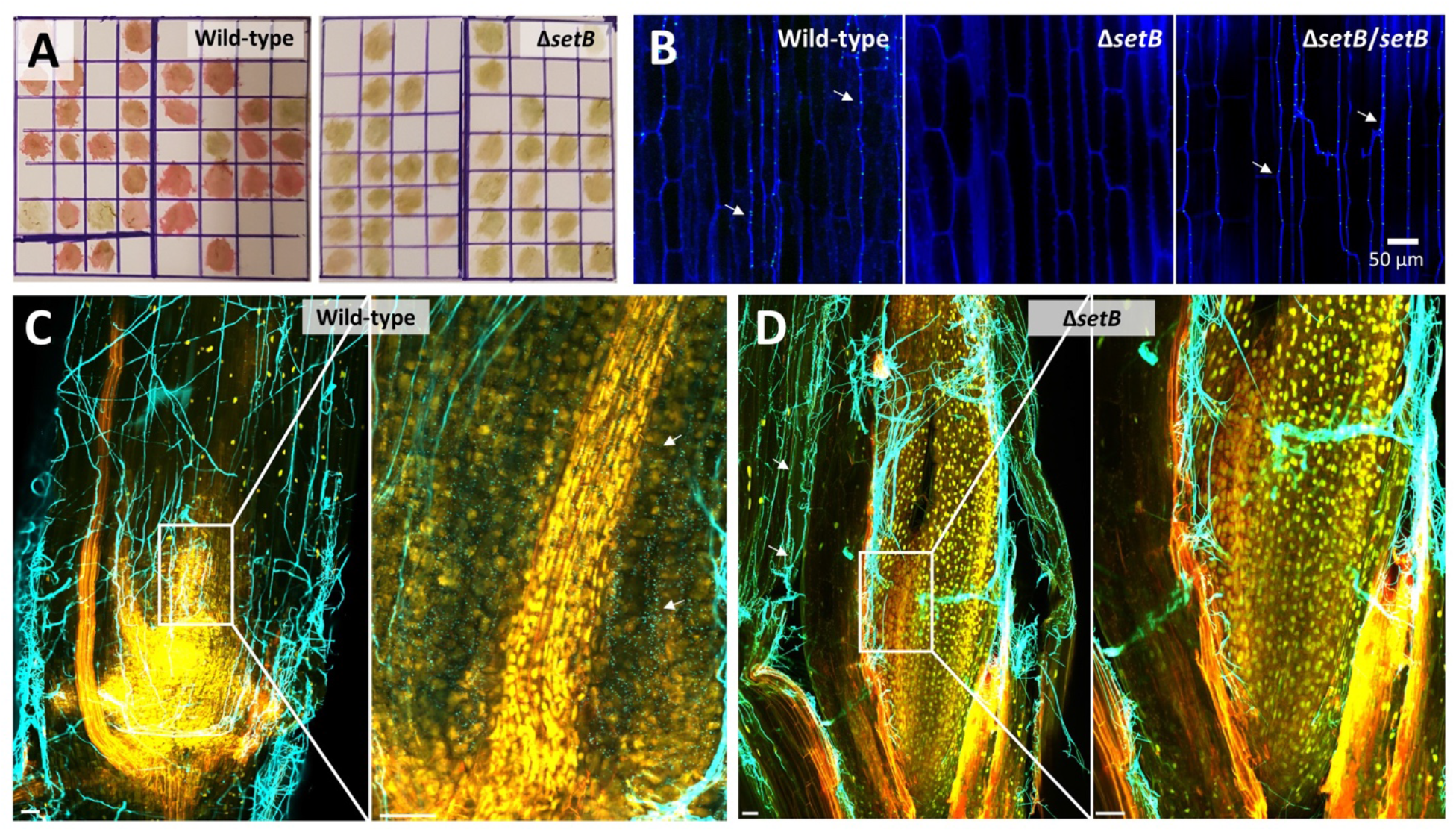
*E. festucae setB* is required for host infection. (A) Immunoblot detection of *E. festucae* in plant tillers, visualized with Fast Red. (B) Confocal microscopy analysis of mature plant tillers. Aniline blue was used to stain cell-wall glucans in hyphae (blue) and WGA-AF488 labels chitin in fungal septa (cyan). Image of Δ*setB* showing absence of hyphae is representative of more than 10 plants analysed. z = 5 μm. (C & D) Confocal microscopy of 2 wpi seedlings stained with WGA-AF488 (in cyan) indicate a presence of endophytic hyphae in seedlings infected with wild-type (C) but not Δ*setB* (D). WGA-AF488 stains chitin in the cell wall of epiphytic hyphae but only stains the septa of endophytic hyphae. Images were generated by maximum intensity projection of z-stacks. Bars = 50 μm.

Subsequently, we sought to determine whether the phenotypes observed for Δ*setB* were caused by the lack of H3K36 trimethylation in this strain. Substitution of a conserved cysteine residue for alanine in the SET domain of *Neurospora crassa* SET-2 has previously been shown to result in a severe reduction in methyltransferase activity, confirming the importance of this amino acid for enzyme activity [11]. We therefore substituted alanine for the corresponding C254 residue in *E. festucae* SetB (Fig 1B) and tested if this allele was able to complement the H3K36 methylation, culture growth and host infection defects of Δ*setB*. Introduction of this *setB*^C254A^ allele only partially rescued the H3K36me3 defect of Δ*setB*, as well as the aberrant accumulation of H3K36me1/2 observed for this strain (Fig 1C). Likewise, the colony growth (Fig 1D) and host infection phenotypes (Table 1) were only partially rescued by introduction of this allele. Although the Δ*setB*/*setB*^*C254A*^ strain was able to infect the host, infection was at a significantly lower rate than the Δ*setB*/*setB* complement or the wild-type strains (Table 1). Plants that were successfully infected with the Δ*setB*/*setB*^*C254A*^ strain were phenotypically similar to wild-type infected plants (S2 Fig). These results suggest that the phenotypes observed for Δ*setB* were due to the H3K36 methylation defects.

### The H3K9 methyltransferase activity of ClrD is also required for host infection by *E. festucae*

The *E. festucae* Δ*clrD* mutant, which is defective in H3K9 mono-, di- and tri-methylation, also has a non-infection phenotype [8, 10]. However, that analysis did not differentiate lack of infection from lack of persistence as host plants were only examined for infection at 10-12 wpi. To discriminate between these two possibilities, we examined seedlings infected with mutant or wild-type at one- and two- wpi using CLSM. At both timepoints epiphytic and endophytic hyphae were observed for wild-type infected seedlings, but only epiphytic hyphae were observed at the inoculation site for Δ*clrD* (Fig 3). These observations suggest that Δ*clrD*, like Δ*setB*, is completely incapable of infecting the host plant. It is interesting to note that in addition to this non-infection phenotype, Δ*clrD* also shares several other phenotypes with Δ*setB*, including a slow growth rate on PDA (Fig 4A) and aberrant hyphal and cellular morphologies (Fig 4B).

**Fig 3.**
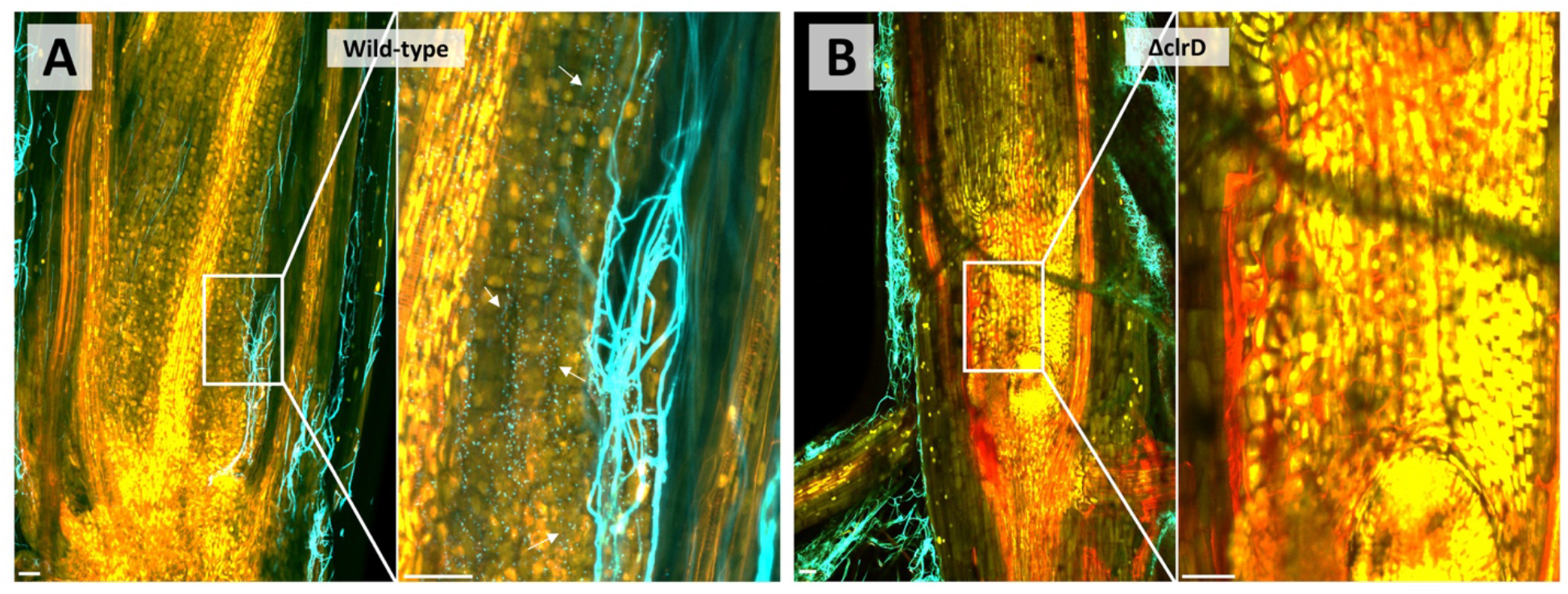
*E. festucae clrD* is required for host infection. Confocal microscopy of 2 wpi seedlings stained with WGA-AF488 (cyan) showing presence of epiphytic and endophytic hyphae in seedlings inoculated with wild-type (A) but only epiphytic hyphae for Δ*clrD* (B). Images were generated by maximum intensity projection of z-stacks. Bars = 50 μm.

**Fig 4.**
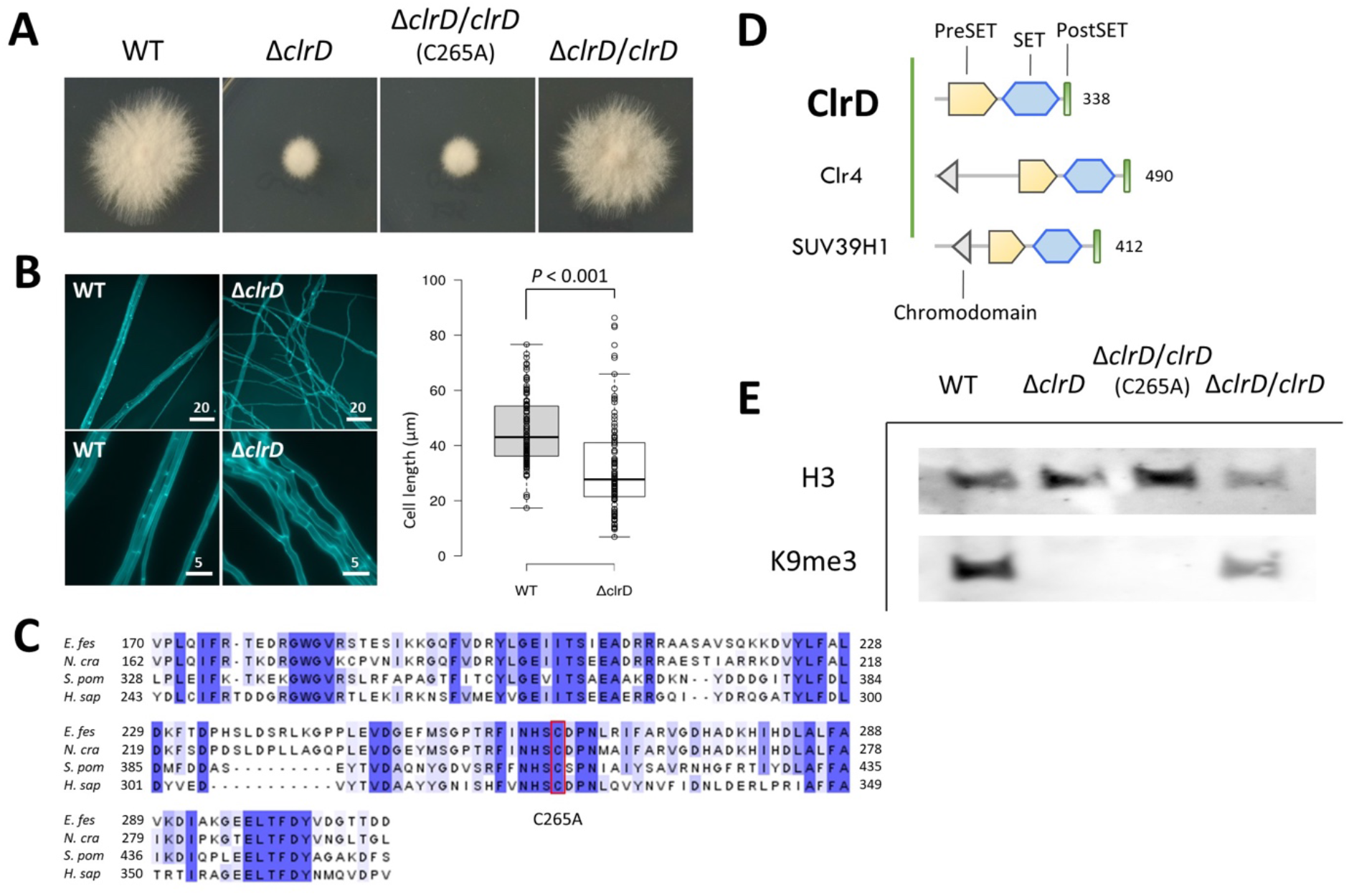
*E. festucae* ClrD is a H3K9 methyltransferase. (A) Colony morphology after 6 days of growth in culture. (B) Deletion of *clrD* affected hyphal morphology and length. Numbers above scale bars refer to length in μm. Boxplots represent more than 80 measurements of hyphal compartment lengths for each strain. The horizontal centre line of the boxes represent the medians and the top and bottom edges of the boxes represent the 75^th^ and 25^th^ percentiles, respectively. (C) Amino acid alignment of SET domains of ClrD and other KMT1 proteins showing conservation of the cysteine 265 residue that was mutated to alanine in this study. (D) Domain structure of *E. festucae* ClrD and the yeast and mammalian orthologs. Numbers indicate protein length in amino acids. (E) Western blot of total histones show reduced H3K9me3 in Δ*clrD*. Relative histone methylation was quantified by comparison to H3 band intensities, with the wild-type value arbitrarily set to 1.

To determine if the lack of H3K9 methyltransferase activity in the Δ*clrD* strain was responsible for the non-infection phenotype of the mutant [8], we tested if a *clrD*^C265A^ allele (Fig 4C & D) could complement the growth and H3K9 methylation defects of the Δ*clrD* mutant. The corresponding C265 residue in KMT1 (metazoan homolog of ClrD) is essential for the histone methyltransferase activity of the protein [27]. Likewise, we found that this allele failed to rescue the Δ*clrD* defects in H3K9 trimethylation (Fig 4E), colony growth (Fig 4A), and host infection (Table 1), indicating that the Δ*clrD* phenotypes observed were due to the absence of H3K9 methylation.

### The infection-negative phenotypes of Δ*setB* and Δ*clrD* are not due to an altered host defense response

We next tested whether Δ*setB* and Δ*clrD* triggered an aberrant host defense response by measuring the expression of host defense genes in seedlings challenged with the mutants. *L. perenne* genes encoding proteins involved in the biotic stress response that were differentially expressed in response to infection with wild-type *E. festucae* Fl1 at seven wpi have previously been reported [28]. From this dataset we selected three of the most highly upregulated genes (m.131905, m.302781 and m.11574) and three of the most highly downregulated genes (m.41989, m.228647 and m.73716) with putative host defense functions (S3 Fig) and analysed their expression in seedlings at five days post inoculation (dpi) with wild-type, Δ*setB*, Δ*clrD*, or in mock-inoculated seedlings. None of these genes were significantly upregulated in plants inoculated with Δ*setB* or Δ*clrD* compared to wild-type (S3 Fig). One gene, m.11574 was significantly downregulated in the Δ*setB* compared to wild-type infected plants. This m.11574 gene was also the only differentially regulated gene in wild-type vs. mock-inoculated plants at five dpi, suggesting that expression of these genes at this early time point is very different to expression in mature plants [28]. These results suggest that the mutants do not induce host defense responses more strongly than the wild-type strain. A host defense response can also present as a brown discoloration of the host tissues surrounding the inoculation site [29, 30]. However, no such browning of the host tissue was observed in seedlings inoculated with either Δ*setB* or Δ*clrD*.

### Δ*clrD* and Δ*setB* are able to infect *L. perenne* if co-inoculated with the wild-type strain but give rise to incompatible associations

Finally we considered the possibility that Δ*clrD* and Δ*setB* do not produce permissive factors for infection, such as small secreted effector proteins. To this end, we tested if the wild-type strain, which would express such factors could rescue infection of the mutants. Seedlings were co-inoculated with mutant and wild-type mycelia at a 4:1 (mutant to wild-type) ratio to maximize the probability of the mutant entering the host. Infection was determined in mature plants (8 wpi) by immunoblot analysis and PCR was subsequently used to distinguish between wild-type and mutant endophyte strains present in these plants. Surprisingly, we were able to detect presence of the Δ*setB* mutant in 5/24 plants (S4A Fig), and the Δ*clrD* mutant in 1/13 plants (S4C Fig). The presence of PCR products for *setB* and *clrD* showed that the wild-type strain was also present in each of these plants (S4B and S4D Fig). This result indicates the wild-type strain facilitates host infection by the Δ*setB* and Δ*clrD* mutants.

To substantiate the above PCR results, we repeated the co-inoculation study using Δ*clrD* or Δ*setB* strains, constitutively expressing eGFP, together with wild-type expressing mCherry. We were able to observe both eGFP- and mCherry-expressing hyphae in the mature plant tissues by CLSM, confirming that both Δ*clrD* and Δ*setB* mutants are able to infect if co-inoculated with the wild-type strain (Fig 5). In addition to individual eGFP- or mCherry-expressing hyphae, we also observed hyphae expressing both eGFP and mCherry (Fig 5A), the result of anastomosis between mutant and wild-type hyphae. A hyphal fusion test in axenic culture showed that the Δ*setB* and Δ*clrD* mutants were indeed capable of fusing with the wild-type strain (S5 Fig). However, the presence of individual eGFP-labelled hyphae in the plant indicates that this fusion with wild-type hyphae is not required for the mutants to enter the host, as it is not possible for anastomosed hyphae to re-segregate. Taken together, these results support the hypothesis that Δ*setB* and Δ*clrD* lack permissive factors for infection, which can be supplied by co-inoculation with the wild-type strain.

**Fig 5.**
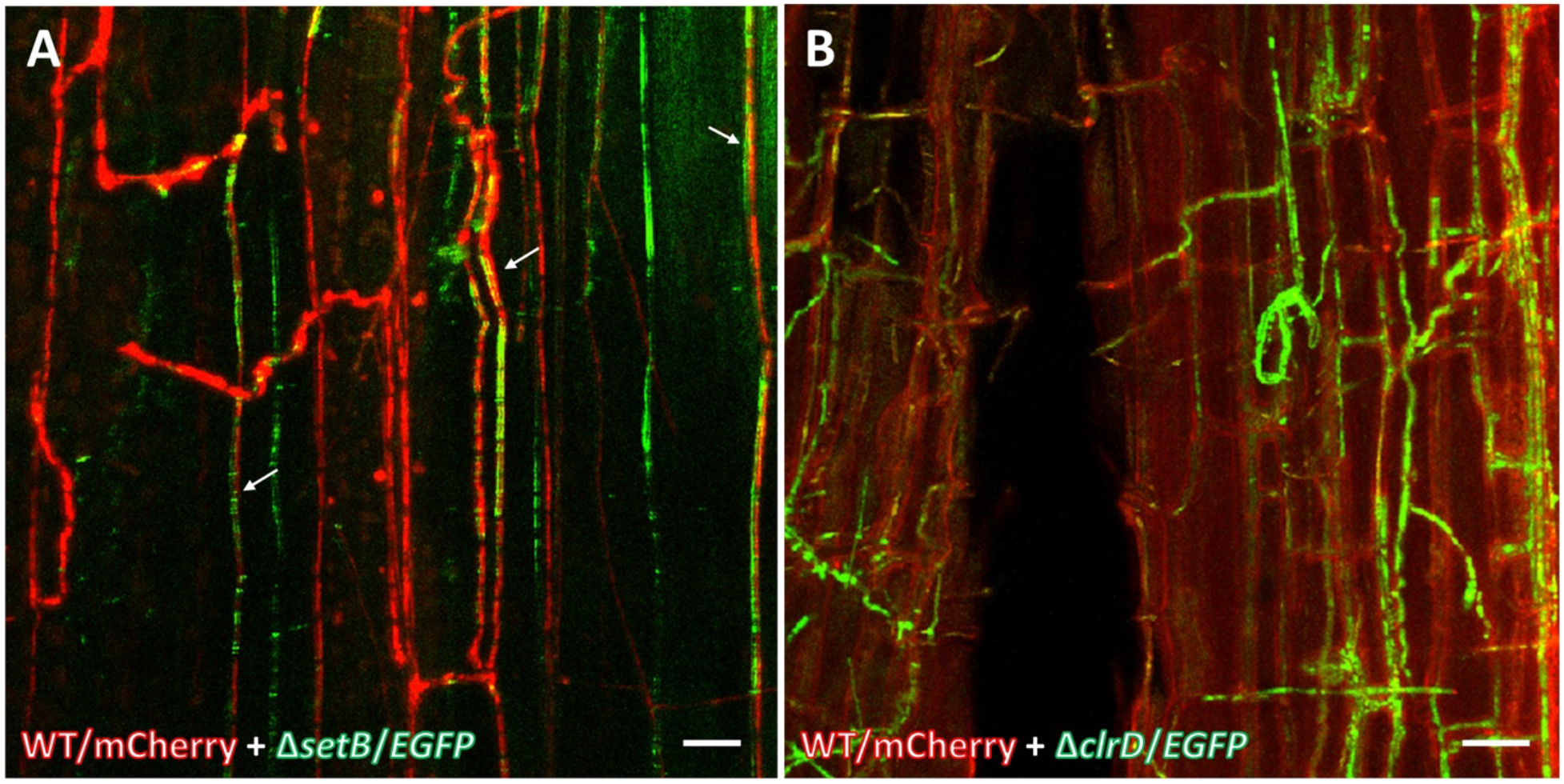
Co-inoculation of Δ*setB* or Δ*clrD* with the wild-type strain enabled infection mutants to colonize the host. (A) Confocal microscopy of plant tillers coinfected with mCherry-tagged wild-type and eGFP-tagged Δ*setB* strains. (B) Confocal microscopy of plant tillers coinfected with mCherry-tagged wild-type and eGFP-tagged Δ*clrD* strains. White arrows indicate hyphae expressing both mCherry and eGFP. Bars = 20 μm.

In addition to performing an immunoblot assay using anti-*E. festucae* antibody to determine infection in the co-inoculation experiment described above, we also performed a replica blot using anti-GFP antibody to determine the presence of the eGFP-labelled mutants in the infected plants. This revealed strong eGFP presence in a number of plants, all of which had underdeveloped root systems and just a single <10 cm tiller, and showed signs of senescence at 8 wpi (S6 Fig). Similar-aged ryegrass plants had >3 tillers and a height of >30 cm, indicating that presence of the Δ*setB* or Δ*clrD* mutants in the host may lead to host stunting. The remaining plants, which did not test positive for eGFP presence in the immunoblot, showed a range of host interaction phenotypes from severe stunting to normal growth (S7A and S7C Fig). To test the hypothesis that the stunted plants contained more mutant hyphae compared to the non-stunted plants, we isolated gDNA from these plants and performed qPCR analysis to measure the relative abundance of *eGFP* (marker) to *pacC* (control), a single-copy gene in *E. festucae* [31] In the Δ*setB*/*eGFP* co-inoculated plants, we detected the *eGFP* gene in four out of five stunted plants (80%), and in one of the six non-stunted plants (17%) (S7B Fig). In the Δ*clrD*/*eGFP* co-inoculated plants, we detected the *eGFP* gene in 11 out of the 14 stunted plants (79%), and in two of the seven non-stunted plants (29%) (S7D Fig). These results indicate that host stunting correlates with presence of the mutants inside the plants. Taken together, these results suggest that while Δ*setB* or Δ*clrD* hyphae are not able to infect the host on their own, when their infection is facilitated by co-inoculation with the wild-type strain, these mutants trigger a host response that results in stunting and premature death.

### Transcriptomics of Δ*clrD* and Δ*setB* provide insights into *E. festucae* host infection process

To determine how the *E. festucae* and *L. perenne* transcriptomes are altered in the early stages of infection with wild-type or mutant strains, high throughput mRNA sequencing was performed on three biological replicates of *L. perenne* seedlings at three dpi with wild-type, Δ*clrD*, Δ*setB*, or a mock inoculation control. Fungal reads were mapped to the most recent *E. festucae* strain Fl1 gene model set [32], and the minimum definition for a differentially expressed gene (DEG) was defined as a statistically significant (S value ≤ 0.005) twofold change in gene expression between conditions (S1 File). This identified 1,027 DEGs in Δ*setB* vs. wild-type (746 downregulated and 281 upregulated), 705 DEGs in Δ*clrD* vs wild-type (333 down and 372 up), and a core set of 215 DEGs (160 down and 55 up) that were differentially regulated in both Δ*clrD* vs. wild-type and Δ*setB* vs. wild-type. Genes encoding predicted *E. festucae* effectors [33] are significantly over-represented in this core gene set (Fishers exact test, *p*=1 × 10^−5^) comprising 5.6% (12/215) of this set compared to 1.2% (99/7,938) of all genes.

Four genes from these DEG sets stood out as candidate infection factors that might contribute to host colonization. These were identified by applying two criteria: first we looked for small secreted protein (SSP)-encoding genes that were upregulated in wild-type *in planta* at three dpi (this study) relative to axenic culture [9], which suggests they may play important roles during host infection (S2 File). Secondly, we searched for genes exhibiting strong differential expression in one or both of the Δ*clrD* and Δ*setB* mutants vs. wild-type. The first gene we identified, named *crbA* (EfM3.044610), encodes a putative 162 aa putative <u>c</u>a<u>r</u>bohydrate-<u>b</u>inding protein, and is downregulated 114-fold to an almost silent state in Δ*setB*. The second gene (EfM3.066990), *dmlA* (<u>d</u>o<u>m</u>ain<u>l</u>ess protein A), while not encoding an SSP, is entirely shut down in Δ*clrD* and encodes an 85 aa protein with no known domains. The third gene, *hybC* (EfM3.007740), encodes an 83 aa putative hydrophobin containing eight cysteine residues, a characteristic feature of hydrophobins [34]. Expression of this gene is upregulated 600-fold at 3 dpi in wild-type vs. culture, and is strongly downregulated in both Δ*setB* (250-fold) and Δ*clrD* (35-fold) vs. wild-type at 3 dpi. The fourth gene, *sspZ* (EfM3.050840), encodes a putative 85 aa small secreted protein. This gene is silent in wild-type axenic culture, minimally expressed at 3 dpi with wild-type, and upregulated in Δ*setB* (7-fold) and Δ*clrD* (61-fold) at 3 dpi. Such upregulated genes are unlikely to encode proteins required for infection but may represent genes that could impair *E. festucae* infection ability if overexpressed. Three of these four genes, *crbA*, *hybC* and *sspZ*, were identified as candidate *E. festucae* effectors [33, 35] (S1 file) and located in the sub-telomeric region of the Fl1 genome In addition *crbA*, *dmlA* and *sspZ* genes are found proximal to AT-rich or repeat elements (S8 Fig), which are regions of the *Epichloë* genome that are particularly enriched for symbiosis genes [32]. Two of these, *crbA* and *sspZ* are subtelomeric (S9 Fig). Interestingly, *crbA* appears to be a component of a four-gene cluster present in some other *Epichloë* species (S1 Table, S10 Fig), which is comprised of a divergently-transcribed gene encoding a chitinase (EfM3.044620), together with genes encoding ankyrin domain protein (EfM3.104630) and a serine/threonine kinase (EfM3.104640) (S10 Fig).

This 3 dpi transcriptome data also provided the opportunity to interrogate the host response to *E. festucae* during the very early stages of the infection. Plant reads were mapped to our *L. perenne* gene model set for which putative functions have been assigned [28], DEG minimum requirements were set as above. Within the category of genes for “biotic” stress (n=830), 30 genes were upregulated and one was downregulated in seedlings inoculated with wild-type compared to mock (S3 File). By comparison, there were no upregulated genes within this category in seedlings inoculated with Δ*setB* or Δ*clrD* compared to the wild-type strain (S4 File). Rather, five genes in Δ*setB* and three in Δ*clrD*, were downregulated compared to the wild-type. The six genes analysed in our RT-qPCR analysis (S4 Fig) were also not differentially expressed in this dataset. These results support the conclusion that Δ*setB* and Δ*clrD* do not induce an aberrant host defense response.

### Functional analysis of genes encoding putative infection proteins

We next generated deletion strains of *hybC*, *crbA* and *dmlA* by targeted homologous recombination, as described above for *setB*, with 48 geneticin resistant transformants screened from each transformation using multiplex PCR. This analysis identified one transformant from each transformation in which the respective target gene was absent (S11A-C Fig). The targeted replacement of these genes was confirmed by PCR amplification across the left and right borders of deletion loci (S11A-C Fig), and multiplex PCRs were repeated with a high cycle number (45 cycles) to confirm absence of each gene in their corresponding mutant strain (S11D Fig). We also generated overexpression strains for *sspZ* by transforming wild-type with a construct placing *sspZ* under the control of the constitutively active P*tefA* promoter (S11E Fig). A total of 24 transformants were screened by qPCR and four transformants that contained the highest construct copy number in the genome, which ranged from 8-10 copies (S11E Fig), were selected for further analysis.

The culture colony morphology of all mutants was similar to the wild-type strain (S12 Fig). To test if the strains were disrupted in their ability to infect the host, they were inoculated into perennial ryegrass seedlings and the infection status of the plants was determined at nine wpi by immunoblotting. All mutants were observed to infect the host (S13 Fig); however, infection rates of the Δ*crbA* mutant, and to a lesser extent the Δ*dmlA* mutant, were significantly lower (S2 Table). There were no differences in the gross morphology of mutant-infected plants from wild-type infected plants at 10 wpi.

## Discussion

The combination of both forward and reverse genetics studies in *Epichloë festucae* have identified key signaling pathways for the symbiotic interaction with the grass host, *Lolium perenne*. These include fungal cell wall integrity [36, 37], pheromone response/invasive growth [38], and stress-activated [39] mitogen-activated protein kinase (MAPK) pathways, cAMP [40], calcineurin [29], pH [31], light [41, 42], reactive oxygen species (ROS) [43–45] and lipid [46] signalling pathways. These genetic analyses uncovered host interaction phenotypes ranging from a mild alteration in the host tiller morphology to severe stunting of tiller growth. In the latter case, mutant hyphae exhibit pathogen-like proliferative growth within the host aerial tissues and colonize the host vascular bundles, phenotypes not observed for the wild-type [47]. *E. festucae* mutants defective in cell-cell fusion all exhibit this antagonistic host interaction phenotype, leading to the hypothesis that hyphal fusion and branching within the host plant are crucial for the establishment of a symbiotic hyphal network [47]. We have also identified a new class of symbiotic mutants defined by deletion of *clrD*, which encodes the histone H3K9 methyltransferase, that completely lack the ability to colonize the host [8]. In this study, we identified another mutant of this class, Δ*setB*, which lacks the histone H3K36 methyltransferase.

As shown by CLSM using aniline blue and WGA-AF488, both Δ*clrD* and Δ*setB* mutants are incapable of infecting and establishing an endophytic network of hyphae within the host, instead forming an epiphytic hyphal network around the site of inoculation. This inability to infect the host leaf cortical tissue is unlikely to be due to the slow growth rate phenotypes of the Δ*clrD* and Δ*setB* mutants because this is a phenotype also observed for the many infection-competent strains of *E. festucae* var *lolii* [48, 49]. *E. festucae* mutants of the small GTPases *racA* and *pakA* (*cla4*), which have similar slow growth phenotypes in culture, are also able to infect perennial ryegrass [50, 51]. In fact, the vigorous epiphytic growth of Δ*clrD* and Δ*setB* and endophytic growth of Δ*racA* and Δ*pakA* suggest that growth in culture is not necessarily a reflection of the growth potential *in planta*.

Although the host and culture phenotypes of both mutants could be fully complemented by the wild-type alleles, the SET domain mutant allele *clrD^C^*^265A^ failed to rescue both the host infection phenotype and the H3K9me3 defect. This result is consistent with the conserved cysteine residue being essential for the catalytic activity of KMT1 [27]. In contrast, the catalytic activity of SetB and the culture and infection phenotypes of the Δ*setB* mutant are only partially dependent on the conserved cysteine 254 residue of the SET domain of the protein. It is likely that the preceding arginine residue (R248), which is highly conserved and crucial for the activity of Set2 in *N. crassa* [11], is additionally required for the methyltransferase activity of SetB. The need for catalytically active ClrD and SetB highlights the important role H3K9 and H3K36 methylation have in regulating *E. festucae* host infection. Given the significance of these marks in transcriptional regulation, it is likely that misregulation of downstream genes are responsible for the infection defects of Δ*clrD* and Δ*setB*.

Surprisingly, co-inoculation with the wild-type strain enabled the mutants to infect and colonize the host plant. This suggests the wild-type strain is able to modulate the host environment to allow infection. However, co-infection resulted in host incompatibility, with host phenotypes varying from mild to severe stunting of the tillers, phenotypes observed previously for some other symbiotic mutants [47]. Severity of the host phenotype correlated with presence of mutant hyphae as determined by qPCR, a result consistent with phenotypes observed for a number of mutants that have a proliferative growth phenotype in the host [43, 45]. The presence of mutant hyphae in the host tissue was also confirmed by CLSM using strains expressing eGFP and mCherry. Importantly, while CLSM showed that fusion between mutant and wild-type hyphae did occur, hyphae with just one fluorescent marker were also present, highlighting that fusion with the wild-type hyphae is not a prerequisite for host colonization by mutant hyphae after co-inoculation.

Since the completion of this study, another *E. festucae* mutant with a non-infection phenotype, Δ*mpkB* has recently been isolated [38]. The *mpkB* gene is a homolog of the *N. crassa* gene *mak-2*, which encodes the MAPK required for pheromone response/invasive growth signaling [52]. As found for *N. crassa* Δ*mak-2* mutants, the *E. festucae* Δ*mpkB* mutant is also defective in cell-cell fusion. The inability of Δ*mpkB* to infect the plant host is unusual as all other *E. festucae* cell-cell fusion mutants isolated to date are still able to colonize host aerial tissues. Given *mpkB* was not differentially expressed in either Δ*clrD* or Δ*setB* mutants at 3 dpi, it is unlikely that the infection defects of these mutants are dependent on MpkB. Similarly, other genes involved in the regulation of hyphal fusion in *E. festucae* are not differentially expressed in the Δ*clrD* and Δ*setB* mutants at 3 dpi, including those of the ROS signaling pathway components encoded by *noxA*, *noxR*, *racA* and *bemA* [43–45], or the cell wall integrity-MAPK pathway components encoded by *mpkA*, *symB* and *symC* [36, 37]. All of these genes, including *mpkB*, are essential for cell-cell fusion in *E. festucae* and for symbiosis with the host; however, the ability of Δ*clrD* and Δ*setB* to anastomose indicates that these are a distinct class of mutants to both Δ*mpkB* and the signaling pathways described above.

Major changes in the host transcriptome occur upon infection with *E. festucae*, suggesting that the endophyte modulates host gene expression to establish a mutualistic symbiotic association [28, 53, 54]. Among the host genes altered in the interaction between *E. festucae* strain Fl1 and *L. perenne* are those involved in biotic stress response, including plant defense genes [28]. However, none of these putative defense genes were aberrantly induced upon inoculation of *L. perenne* seedlings with the Δ*clrD* or Δ*setB* strains at 3 dpi. RT-qPCR analysis for six of these putative defense genes that were previously shown to be differentially expressed at seven wpi [28] showed no statistically significant change in expression in host tissues from the infection site of seedlings at five dpi with the exception of one (m.11574), encoding a leucine-rich repeat receptor-like protein, which was instead downregulated in Δ*setB*-inoculated plants. This result was confirmed by a full transcriptome analysis of differences in host gene expression at three dpi where just five putative host defense genes were differentially expressed, all of which were downregulated in the Δ*clrD* or Δ*setB*-inoculated seedlings compared to seedlings inoculated with wild-type. In addition, the absence of any visual necrotic response as observed for ryegrass seedlings inoculated with a calcineurin mutant [29] suggests a hypersensitive response is not responsible for the lack of colonization by Δ*clrD* and Δ*setB*. Therefore, these mutants appear to be inherently incapable of infecting the host. In this respect, the ability of the wild-type strain to rescue infection in the co-inoculation experiments suggests that the wild-type strain secretes some factor into the extracellular environment which allows for host infection by these mutants.

Many plant-pathogenic fungi secrete small proteins that play important roles in pathogenicity and virulence. The tomato pathogen *Cladosporium fulvum* secretes LysM domain-containing effectors Avr4 and Ecp6 which function to bind and prevent host recognition of chitin, a major component of the fungal cell wall and a potent immunogen in plants [55, 56]. Other effectors directly modulate the host plant defense response, such as the *Ustilago maydis* effector Pit2 which inhibits host papain-like cysteine proteases central to the host apoplastic immunity [57, 58]. However, fungal effectors can also be important in mutualistic interactions as observed in the ectomycorrhizal fungus *Laccaria bicolor* which secretes an effector, MiSSP7, that regulates the symbiosis by modulating the host jasmonic acid signaling pathway [59, 60]. Modulation of the host response by wild type secretion of effectors is one possible explanation for why co-inoculation of wild-type together with either Δ*clrD* or Δ*setB* enables these strains to infect the host. Candidate infection factors therefore include predicted small secreted proteins (SSPs) that are encoded by genes exhibiting both high expression by the wild-type *in planta*, and low expression by the Δ*clrD* and/or Δ*setB* mutants *in planta*. From the four predicted SSP genes selected for functional analysis, deletion of *crbA*, and to a lesser extent *dmlA*, led to significantly reduced infection rates. Deletion of *hybC* and overexpression of *sspZ*, which was upregulated in both Δ*clrD* and Δ*setB*, had no effect on the symbiotic interaction phenotype of *E. festucae*, possibly due to functional redundancy between SSPs [33, 61, 62]. These results show that while the ability to infect the host is a function of the cumulative effects of multiple gene products, some small secreted proteins can, and do, confer a significant individual contribution towards the efficiency of this process. Interestingly, *crbA* is also downregulated in the *E. festucae* Δ*hepA* mutant, which lacks the H3K9me3-binding Heterochromatin Protein I [9], as well as in three other symbiotic mutants Δ*sakA*, Δ*noxA* and Δ*proA* [63]. Intriguingly, *crbA* appears to be a component of a cluster of four genes encoding proteins important for cell wall (chitinase) and cell membrane (ankyrin) modifications. With the exception of *hybC* the three other genes analysed here are all found close to AT-rich or repeat-rich blocks of the genome, regions particularly enriched for fungal effectors [64, 65] and symbiotic genes [32].

Given the observation that chitin is masked or modified in endophytic but not epiphytic or axenic hyphae of *E. festucae* [66], remodelling of the fungal cell wall is also likely to be important for colonization and symbiosis. However, it is difficult to see how the wild-type strain could act *in trans* to complement this defect in the mutants. Instead, incomplete cell wall remodelling may explain the host incompatibility response elicited by Δ*clrD* and Δ*setB* once inside the plant. In addition, while the Δ*crbA* and Δ*dmlA* mutants are compromised in their ability to infect, plants that are infected with these mutants do not reproduce the incompatibility phenotypes observed for plants co-infected with wild-type and Δ*clrD* or Δ*setB*. This is not surprising given that H3K9 and H3K36 methylation defects would affect the expression of a large number of genes, resulting in more dramatic phenotypes for these mutants. However, these host colonization and symbiosis defects appear to be specific to these two H3 marks as mutations that affect H3K4 and H3K27 methylation have little or no impact on the host interaction phenotype [8, 10].

H3K9 and H3K36 methylation play important roles in regulating fungal development and pathogenicity. Silencing of the *clrD* homolog, *DIM-5*, in *Leptosphaeria maculans* led to the aberrant overexpression of effector genes located near AT-isochores and attenuated pathogenicity in oilseed rape [67]. Deletion of the *clrD* and *setB* homologs, *Mokmt1* and *Mokmt3*, reduced the pathogenicity of *Magnaporthe oryzae* across several host plants [20]. Similarly, deletion of *set2* in *Fusarium verticillioides* led to reduced virulence and production of the mycotoxin bikaverin [21]. The growth defects of the Δ*clrD* and Δ*setB* mutants observed in this study are consistent with those observed in the *N. crassa* and *Aspergillus nidulans* mutants for these genes [22–25]. Interestingly, deletion of the *set2* homolog *ash1* in *Fusarium fujikuroi* led to a more severe developmental phenotype than deletion of *set2*, which is characterised by instability of subtelomeric chromosome regions and loss of accessory chromosomes, phenotypes associated with the proposed role of this paralog in DNA repair. Both mutants also had reduced host pathogenicity [19]. Given the global roles of ClrD and SetB in the maintenance of H3K9 and H3K36 methylation in the genome, it is likely that these proteins are not directly responsible for the infection ability of *E. festucae*, but rather this role is performed by other genes under their regulation.

In conclusion, we show here that ClrD-catalysed H3K9 and SetB-catalysed H3K36 methylation are crucial in regulating the ability of a fungal symbiont to infect its host. The results of this study also underscore the importance of further analysis into the symbiotic roles and mode of action of the small secreted protein encoded by the *crbA* gene, which appears to be important for *E. festucae* infection efficiency.

## Materials and methods

### Fungal growth conditions, transformation and inoculation

Bacterial and fungal strains, plasmids, and plant material used in this study are listed in S3 Table. *E. festucae* strains were grown at 22°C on 2.4% (w/v) potato dextrose agar or broth with shaking at 200 rpm. *E. festucae* protoplasts were prepared and transformed as previously described [68, 69]. Inoculation of *E. festucae* perennial ryegrass seedlings was performed as previously described [70]. Plants were maintained in root trainers in an environmentally controlled growth room at 22°C with a photoperiod of 16 h of light (approximately 100 μE/m^2^/s), and presence of endophyte was detected by immunoblot using anti-*E. festucae* antibody (AgResearch, Ltd), or anti-GFP antibody (Abcam ab290) as previously described [71]

### Generation of DNA constructs and mutants

A list of PCR primers used in this study is provided in (S4 Table). Plasmid pYL23, which contains the *setB* replacement construct, was generated by Gibson assembly [72] from DNA fragments containing the 5’ and 3’ regions flanking *setB* that were amplified from an *E. festucae* Fl1 genomic DNA template using primer pairs YL286F/R and YL288F/R, respectively; a P*trpC*-*nptII*-T*trpC* gene expression cassette that confers geneticin resistance that was amplified from pSF17.1 using primers YL287F/R; and a *Nde*I-linearised pUC19 vector sequence. A linear fragment was excised from pYL23 by *Pac*I/*Spe*I digestion and used for transformation of wild-type *E. festucae* strain Fl1 protoplasts to generate Δ*setB* strains. A 4.3 kb DNA fragment covering the *setB* gene including promoter and terminator sequences was amplified from wild-type *E. festucae* genomic DNA using primers YL330F/R and ligated by blunt-end cloning into *Sna*BI-linearised pDB48 to generate the *setB* complementation plasmid pYL30. Plasmid pYL31, containing the *setB*^C254A^ gene, was generated by site-directed mutagenesis of pYL30 using primer pair YL378F/R. Plasmid pYL32, containing the *clrD*^C265A^ gene, was generated by site-directed mutagenesis of pTC40 using primers YL379F/R. The sequence fidelity of all plasmid inserts were confirmed by sequencing. For fluorescent tagging studies, Δ*setB* and Δ*clrD* protoplasts were transformed with pCT74 harboring *eGFP* under the control of the *toxA* promoter.

Plasmid pYL41 containing the *hybC* replacement construct was generated by Gibson assembly from DNA fragments amplified from an *E. festucae* Fl1 genomic DNA template using primers YL459F/R (*hybC* 5’ flank) and YL461aF/R (*hybC* 3’ flank); the P*trpC*-*nptII*-T*trpC* cassette amplified from pSF17.1 using primers YL460F/R, and *Nde*I-linearised pUC19. A linear fragment for transformation was amplified from pYL41 by PCR using primers YL467F/R. The *crbA* replacement construct-containing plasmid pYL43 was similarly generated from DNA fragments amplified using primers YL464F/R (5’ flank), YL466F/R (3’ flank), YL465F/R (*nptII* cassette), and *Nde*I-linearised pUC19. A linear fragment for transformation was amplified from pYL43 by PCR using primers YL475F/R. The *dmlA* replacement construct-containing plasmid pYL44 was similarly generated from DNA fragments amplified using primer YL486F/R (5’ flank), YL488F/R (3’ flank), YL487F/R (*nptII* cassette), and *Nde*I-linearised pUC19. A linear fragment for transformation was amplified from pYL44 by PCR using primers YL486F/488R. The *sspZ* overexpression constructs pYL45 (with FLAG) and pYL46 (native) were generated by Gibson assembly. For pYL45, the assembled DNA fragments included P*tef* amplified from pYL3 using primers YL496F/R, *sspZ* amplified from wild-type *E. festucae* full-length cDNA using primers YL497F/R, T*tub* amplified from pNR1 using primers YL498F/335R, and *Sna*BI-linearised pDB48. For pYL46, the assembled DNA fragments included P*tef* amplified with primers YL496F/496Rb from pYL3, *sspZ* was amplified from wild-type *E. festucae* full-length cDNA using primers YL497Fb/497R, T*tub* amplified from pNR1 using primers YL498F/335R, and *Sna*BI-linearised pDB48.

For Southern blot analysis [73], DNA was digested and separated by electrophoresis, transferred to positively charged nylon membrane (Roche) and fixed by UV light cross-linking in a Cex-800 UV light cross-linker (Ultra-Lum) at 254 nm for 2 min. Labelling of DNA probes, hybridization, and visualization were performed using the DIG High Prime DNA Labeling & Detection Starter Kit I (Roche) as per the manufacturer’s instructions.

### RNA isolation, reverse transcription and quantitative PCR

Fungal and plant tissue were homogenized with mortar and pestle in liquid nitrogen and RNA was isolated using TRIzol (Invitrogen). For RT-qPCR analysis, cDNA was synthesized using the QuantiTect Reverse Transcription Kit (Qiagen) as per the manufacturer’s instructions. qPCR was performed using the SsoFast™ EvaGreen Supermix (Bio-Rad) on a LightCycler® 480 System (Roche) according to the manufacturer’s instructions with two technical replicates per sample. RT-qPCR was performed using absolute quantification and target transcript levels were normalized against the *E. festucae* reference genes *S22* (ribosomal protein S22; EfM3.016650) and *EF-2* (elongation factor 2; EfM3.021210) [8] or the *L. perenne* reference genes *POL1* (RNA polymerase I; m.40164) and *IMPA* (importin-a; m.15410) [28]. In all cases similar results were obtained by normalizing with either gene and only results normalized with *S22* or *POL1* are presented.

### Histone extraction and western blotting

Histone extraction and western blot were performed as previously described [10]. In brief, fungal tissues were ground to a fine powder in liquid nitrogen, and nuclei were isolated by glycerol gradient centrifugation, sonicated, and histones were subsequently isolated by acid extraction.

### Microscopy and hyphal fusion

Growth and morphology of hyphae *in planta* was determined by staining leaves with aniline blue diammonium salt (Sigma) to stain fungal ß-1,3-glucans and Wheat Germ Agglutinin conjugated-AlexaFluor488 (WGA-AF488; Molecular Probes/Invitrogen) to stain chitin, as previously described [66]. While aniline blue itself is not fluorescent, there is a minor fluorochrome component present, Sirofluor, that is fluorescent [74]. Hyphal growth and fungal cellular phenotypes were documented by CLSM using a Leica SP5 DM6000B (Leica Microsystems) confocal microscope outfitted with a 10×, NA 0.4, 40×, NA 1.3 or 63× NA 1.4 oil immersion objective lens. WGA-AF488, to detect chitin, and aniline blue, to detect β-1,3-glucan, were excited at 488 nm and 561 nm, respectively, and their emission spectra collected at 498-551 nm and 571-632 nm respectively.

The culture eGFP-mRFP cell-cell fusion assays, were performed as previously described [36]. For the coinfected mCherry-tagged wild-type and eGFP-tagged Δ*setB* strains and mCherry-tagged wild-type and eGFP-tagged Δ*clrD* strains, unfixed pseudostem samples were examined by CLSM (Leica SP5 DM6000B (Leica Microsystems)) using 488 nm and 561 nm DPSS laser.

### Bioinformatic analysis and transcriptomics

The genome sequences of *E. festucae* were retrieved from the *E. festucae* Genome Project database hosted by the University of Kentucky (http://csbio-l.csr.uky.edu/endophyte/cpindex.php) [26]. Protein domains were analysed with the SMART web-based tool (http://smart.embl-heidelberg.de/) [75, 76]. Multiple amino acid sequence alignments were generated with Clustal Omega (http://www.ebi.ac.uk/Tools/msa/clustalo/) [77, 78].

Scripts for the transcriptome and statistical analyses performed in this study are available from https://github.com/klee8/histone_mutants. Here we describe briefly each step of the analyses. The gene expression of Δ*setB* and Δ*clrD* was compared with wild-type by negative binomial regression for samples harvested three dpi. High-throughput mRNA sequencing was performed on three biological replicates for each of the wild-type, mock, Δ*setB* and Δ*clrD* associations. Samples were indexed and pooled before being sequenced across several Illumina NovaSeq 6000 lanes to reduce sample-to-sample technical variation. RNAseq read quality was assessed with FastQC v0.11.8 [79]. Adapter and poor-quality reads were removed with Trimmomatic v0.38 [80].

To find expression levels for each gene, read counts of each sample aligned against the *Epichloë* gene model set [32] were estimated with Salmon v0.13.1 [81]. The count data was imported into R v3.6.0 [82] using tximport v1.10.1 [83]. Genes that were significantly differentially expressed between mutant and wild type (s-value ≤ 0.005 and ≥ log 2-fold change) were identified with R package DESeq2 v1.22.2 [84] using the log2 fold shrinkage estimator implemented in apeglm v1.4.2 [85]. We accounted for multiple testing by using the s-value of Stephens [86].

A core set of highly up- and down-regulated genes for the Δ*setB* and Δ*clrD* mutants were identified using R. Genes were only included if their differential expression was in the same direction (i.e., all upregulated or all downregulated) in both mutants. Genes were annotated for putative signal peptides using SignalP v 5.0 [87, 88] and putative functions for encoded proteins were annotated using the online version of PANNZER2 [89]. The Δ*setB* and Δ*clrD* transcriptome data used here is available from the Sequence Read Archive (SRA) under BioProject PRJNA556310. A list of the individual biosample numbers is provided in S5 Table. Similarly, the hypothesis that genes in Δ*setB* and Δ*clrD* mutants are differentially expressed *in planta* compared to axenic culture was tested by logistic regression, using previously published RNAseq data [9]. Analysis of *L. perenne* gene expression was carried out as for the *Epichloë* mutant gene expression analysis using the same quality-controlled RNAseq data, except that reads were aligned to the previously published *L. perenne* gene model set [28].

## Acknowledgements

We thank Matthew Savoian (Manawatu Microscopy and Imaging Centre, Massey University) for technical advice and Daniel Berry and Berit Hassing (Massey University) for comments on this manuscript.

## Supporting Information

**S1 Fig. Strategy for deletion of *E. festucae* setB and confirmation by Southern hybridisation analysis and PCR.**

**S2 Fig. Phenotype of plants infected with Δ*setB*/*setB*^(C254A)^ strain. Plants were photographed at 8 wpi.**

**S3 Fig. Expression of host defense genes in *ΔsetB*- and *ΔclrD*- inoculated plants.**

**S4 Fig. Co-inoculation with the wild-type strain allowed host infection by Δ*setB* and Δ*clrD*.**

**S5 Fig. Hyphal fusion ability is not affected in Δ*setB* and *ΔclrD*.**

**S6 Fig. Immunoblot and phenotype of co-inoculated plants at 8 wpi.**

**S7 Fig. Host stunting correlates with the presence of mutant hyphae in co-inoculated plants.**

**S8 Fig. Candidate infection genes are located near AT-rich or repeat-rich regions.**

**S9 Fig. Chromosome location of four candidate infection genes in *E. festucae* strain Fl1.**

**S10 Fig. Microsyntenic comparison of the 10 kb *crbA* loci in *Epichloë* species possessing *crbA* orthologs.**

**S11 Fig. Strategy for deletion of *E. festucae crbA*, *dmlA*, *hybC* and overexpression of *sspZ*.**

**S12 Fig. Culture phenotype of *E. festucae* wild-type, *crbA*, *dmlA*, *hybC* and *sspZ* mutants on PDA.**

**S13 Fig. Plant infection phenotype for *E. festucae crbA*, *dmlA*, *hybC* and *sspZ* mutants.**

**File S1. Differences in gene expression between *E. festucae* WT and either Δ*clrD* or Δs*etB* at 3 dpi in *L. perenne*.**

**File S2. Differences in gene expression between** *E. festucae* **WT in planta at 3 dpi versus in axenic culture.**

**File S3. Differences in gene expression of *L. perenne* infected with *E. festucae* WT and mock infected.**

**File S4. Differences in gene expression of *L. perenne* infected with *E. festucae* WT and either Δ*clrD* or Δs*etB* at 3 dpi.**

**Table S1: Homologues of *crbA* and *dmlA* in other *Epichloe* species.**

**Table S2: Host infection rates of *E. festucae crbA*, *dmlA*, *hybC* and *sspZ* mutants.**

**Table S3: Biological material.**

**Table S4: Primers used in this study.**

**Table S5: Biosample numbers for transcriptome data.**

## Author Contributions

**Conceptualization:** Yonathan Lukito, Tetsuya Chuo, David J. Winter, Murray P. Cox, Barry Scott.

**Data curation:** Kate Lee, David J. Winter.

**Formal analysis:** Yonathan Lukito, Kate Lee, Nazanin Noorifar, Kimberly A. Green, David J. Winter, Tracy K. Hale, Tetsuya Chuo, Linda J. Johnson, Murray P. Cox, Barry Scott

**Funding acquisition:** Linda J. Johnson, Murray P. Cox, Barry Scott

**Investigation:** Yonathan Lukito, Kate Lee, Nazanin Noorifar, Kimberly A. Green, David J. Winter, Arvina Ram,

**Project administration:** Tracy K. Hale, Tetsuya Chuo, Linda J. Johnson, Murray P. Cox, Barry Scott

**Resources:** Linda J. Johnson, Murray P. Cox, Barry Scott

**Software:** Kate Lee, David J. Winter, Murray P. Cox,

**Supervision:** Tracy K. Hale, Tetsuya Chuo, Linda J. Johnson, Murray P. Cox, Barry Scott

**Writing:** Yonathan Lukito, Barry Scott

